# Condensins are essential for *Pseudomonas aeruginosa* corneal virulence through their control of phenotypic programs

**DOI:** 10.1101/2020.12.08.416081

**Authors:** Hang Zhao, April L. Clevenger, Phillip S. Coburn, Michelle C. Callegan, Valentin Rybenkov

## Abstract

*Pseudomonas aeruginosa* is a significant opportunistic pathogen responsible for a variety of human infections. Its high pathogenicity resides in a diverse array of virulence factors and an ability to adapt to hostile environments. We report that these factors are tied to the activity of condensins, SMC and MksBEF, which primarily function in structural chromosome maintenance. This study revealed that both proteins are required for *P. aeruginosa* virulence during corneal infection. The reduction in virulence was traced to broad changes in gene expression. Transcriptional signatures of *smc* and *mksB* mutants were largely dissimilar and non-additive, with the double mutant displaying a distinct gene expression profile. Affected regulons included those responsible for lifestyle control, primary metabolism, surface adhesion and biofilm growth, iron and sulfur assimilation, and denitrification. Additionally, numerous virulence factors were affected, including type 3 and type 6 secretion systems, hemagglutinin, pyocin and macroglobulin production, and a host of virulence regulators. *in vitro* properties of condensin mutants mirrored their transcriptional profiles. MksB-deficient cells were impaired in pyocyanin, c-di-GMP production, and sessile growth whereas *smc* mutants mildly upregulated c-di-GMP, secreted fewer proteases and were growth deficient under nutrient-limiting conditions. Moreover, condensin mutants displayed an abnormal regulation upon transition to stationary phase. These data reveal that condensins are integrated into the control of multiple genetic programs related to epigenetic and virulent behavior, establishing condensins as an essential factor in *P. aeruginosa* ocular infections.

**Author Summary:** Bacterial pathogenicity is a complex phenomenon dependent on the ability of a bacterium to thrive in a hostile environment while combating the host using an array of virulence factors. This study reports that pathogenicity is also tied to structural chromosome maintenance through condensins, proteins that are responsible for the global organization of the chromosome. We show that the two *Pseudomonas aeruginosa* condensins, SMC and MksB, act as global regulators of gene expression. The inactivation of SMC and MksB induces opposite regulatory programs in the cell that resemble those observed during the acute and chronic phases of infection. A substantial portion of this regulation is mediated by the intracellular signaling network of *P. aeruginosa*. Accordingly, virulence regulation is altered in condensin mutants. The results were validated by genetic, phenotypic and virulence studies of condensin mutants. Overall, these data establish condensins as an essential factor during ocular *P. aeruginosa* infections revealing their involvement in the regulatory virulence network and the control of the bacterial lifestyle.

## Introduction

*Pseudomonas aeruginosa* is one of the primary causes of bacterial keratitis that often leads to severe eye damage and vision loss (1-3). The high pathogenicity of *P. aeruginosa* has been attributed to its adaptability to diverse conditions (4), intrinsically high resistance to antibiotics (5), its secretion of numerous virulence factors, and a severe inflammatory response (6-8). The emergence and spread of multidrug resistance further reduces the efficiency of existing treatments and necessitates the search for new therapeutic targets (9).

Recently, condensins were identified as a novel factor essential for *P. aeruginosa* pathogenicity (10). Condensins are large multisubunit protein complexes responsible for global chromosome organization (11-13). These proteins act as ATP-controlled macromolecular clamps that bring distant DNA segments together, establishing a giant loop chromosome organization (14, 15). *P. aeruginosa* carries two condensins with specialized functions, MksBEF and SMC (16, 17). Unexpectedly, these proteins have also been implicated in the regulation of gene expression and the switching between distinct physiological states of the bacterium (10). SMC deficient cells upregulate exopolysaccharide production and display traits typical for sessile growth. In contrast, inactivation of MksB, whether in the presence or absence of SMC, impairs biofilm formation and facilitates planktonic growth (10). Inactivation of SMC and MksB reduced *P. aeruginosa* pathogenicity in a mouse model of lung infection (10). Curiously, the decline in pathogenicity appeared to correlate with the physiological state of the bacterium rather than the severity of genetic defects since the single *Δsmc* mutant was less virulent than the double mutant *ΔsmcΔmksB*.

We explored here the role of condensins in a mouse model of ocular infection. Deletion of either *smc* or *mksB* accelerated clearance of *P. aeruginosa* from a scratched cornea. This effect was even stronger for the double *ΔsmcΔmksB* mutant. Transcriptomic analysis of condensin mutants revealed that each protein controls the expression of multiple lifestyle and virulence regulons and regulators. Moreover, the transcriptional signature of the double mutant was distinct from both the *smc* and *mksB* strains, suggesting that the bulk of regulation in the mutants involves triggered genetic programs. A phenotypic survey confirmed predictions of the transcriptomic analysis revealing that besides fitness defects, condensin mutants are asymmetrically impaired in the production of virulence factors. Furthermore, condensins showed genetic interactions with several virulence regulators, including *rsmZ*, VreI and BexR, and oppositely affect c-di-GMP production. Overall, these data support the view that inactivation of condensins in *P. aeruginosa* results in a broad dysregulation of gene expression that leads to a reduction in corneal virulence.

## Results

### *Deletion of condensins attenuates virulence of* P. aeruginosa

Infection with PAO1 caused significant inflammation and corneal ulceration, as has been described previously (18-21). Figure 1A represents images taken of the same mouse in each group on three consecutive days. On postinfection day 1, corneas infected with PAO1 showed a diffuse but significant corneal haze consistent with infiltrating inflammatory cells. On postinfection days 2 and 3, corneas infected with PAO1 developed distinct ulcers with denuded corneal epithelium and cellular infiltration into those areas. Corneas infected with the *Δsmc* mutant also developed a diffuse corneal haze on postinfection days 1-3, but the corneal changes in these eyes were not as significant as those observed in eyes infected with PAO1. Corneas infected with the *Δsmc* mutant had notable cellular infiltrates on postinfection day 3, but no corneal ulceration. In stark contrast, eyes infected with the *ΔmksB* mutant or *ΔsmcΔmksB* double mutant showed no signs of corneal infection on postinfection days 1-3.

**Figure 1.**
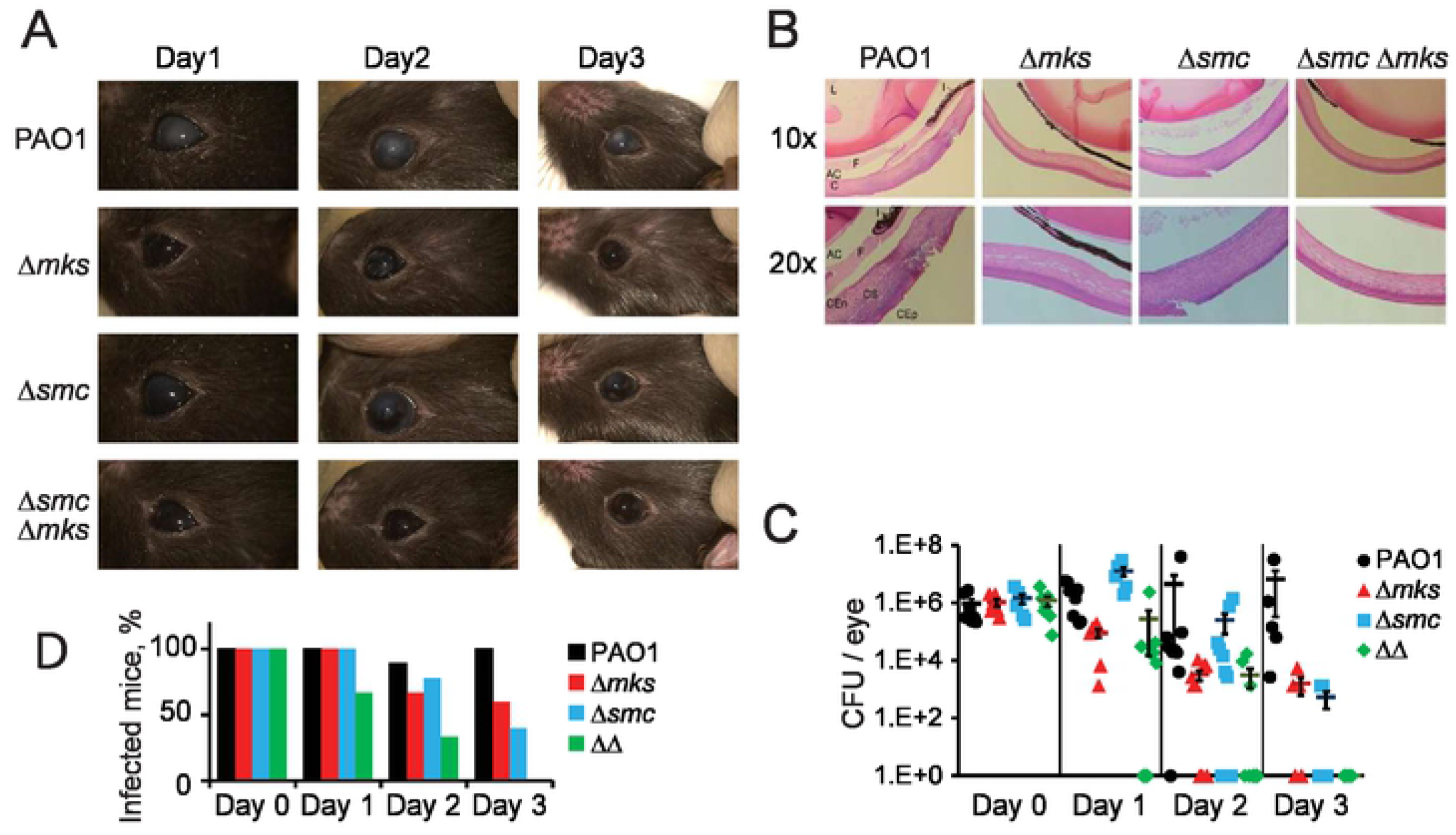
*P. aeruginosa* keratitis. (A) Biomicroscopy on postinfection days 1-3 of eyes topically infected with 10^6^ CFU of *P. aeruginosa* PAO1 or condensin mutants. Images are representative of N=3 eyes/group/time point. (B) Histology of *P. aeruginosa* keratitis. Hematoxylin/eosin histology on postinfection day 1 of eyes topically infected with 10^6^ CFU of *P. aeruginosa* PAO1 or condensin mutants. Images are representative of N=3 eyes/group. Magnification of the objective is indicated on the left. L = lens; I = iris; F = fibrin; AC = anterior chamber; C = cornea; Cen = corneal endothelium; CS = corneal stroma; Cep = corneal epithelium. (C) CFU/eye on postinfection days 1-3. Mean ± SEM for N ⩾ 5 eyes/group. (D) Percent of eyes infected with *P. aeruginosa* PAO1 or condensin mutants (N ⩾ 5 eyes/group).

Figure 1B represents histology of eyes infected with PAO1 or the three condensin mutants on postinfection day 1, when obvious differences in pathology were first noted. Corneas infected with PAO1 demonstrated significant cellular infiltrate into the corneal epithelium and stroma and epithelial damage at the site of infection. Fibrin and immune cell infiltration into the anterior chamber was also observed. Corneas infected with the *Δsmc* mutant also had significant cellular infiltrate into the corneal epithelium and stroma, and epithelial damage at the site of infection. The extent of cellular infiltration in corneas infected with PAO1 or the *Δsmc* mutant appeared similar. In contrast, corneas infected with the *ΔmksB* mutant or the *ΔsmcΔmksB* double mutant appeared normal, with no significant infiltration of immune cells, no damage to the corneal epithelium or stroma, and no fibrin or infiltrating cells in the anterior chamber. The histology data on postinfection day 1 corroborated the biomicroscopic findings, further demonstrating the differences in keratitis pathology caused by the condensin mutant strains. These results indicated that a mutation in MksB, but not SMC, resulted in very minimal pathologic changes in the cornea during keratitis caused by *P. aeruginosa*.

The ocular burden of *P. aeruginosa* PAO1 and its condensin mutants is summarized in Figure 1C. Eyes were infected with approximately 10^6^ CFU of each strain (postinfection day 0, P≥0.29). On postinfection day 1, the PAO1 corneal burden was significantly greater than that of the *ΔmksB* and *ΔsmcΔmksB* mutants (P≤0.0005). The corneal burden of the *Δsmc* mutant was significantly greater than that of all other strains (P≤0.02), including PAO1. On postinfection day 1, all eyes infected with PAO1, and the *ΔmksB* and *Δsmc* mutants had detectable CFU, while 66.7% of eyes infected with the *ΔsmcΔmksB* double mutant had detectable CFU (Fig. 1D). On postinfection day 2, the corneal burden of PAO1 was significantly greater than that of the *ΔmksB* and *ΔsmcΔmksB* double mutant (P≤0.006), but also varied, with the majority of eyes retaining 10^4^ to 10^5^ CFU. The corneal burden of the *Δsmc* mutant was significantly greater than that of the *ΔsmcΔmksB* mutant (P=0.03) but similar to that of PAO1 and the *ΔmksB* mutant (P≥0.09). The corneal burden of eyes infected with the *Δsmc* mutant varied from undetectable to 10^6^ CFU. The corneal burden of eyes infected with the *ΔmksB* and *ΔsmcΔmksB* mutants were similar (P=0.35) and varied from undetectable to 10^4^ CFU. On postinfection day 2, the numbers of eyes with detectable CFU were lower than that observed on postinfection day 1 (PAO1, 88.9%; *ΔmksB*, 66.7%; *Δsmc*, 77.8%; and *ΔsmcΔmksB*, 33.3%). On postinfection day 3, the corneal burden of PAO1 was greater than that of the other groups (P≤0.015) and varied from 10^3^ to 10^7^ CFU. All eyes infected with PAO1 had detectable CFU. The corneal burden of the *ΔmksB* and *Δsmc* mutants was similar on postinfection day 3 (P=0.68). In these two groups, the numbers of eyes with detectable CFU were lower on day 3 than that observed on day 2 (*ΔmksB*, 60%; *Δsmc*, 40%). All eyes infected with the *ΔsmcΔmksB* mutant had no detectable CFU on postinfection day 3. These data suggest that mutating condensins in *P. aeruginosa* altered the fitness of these organisms to survive in the mouse cornea, with the double mutation in *ΔmksB* and *Δsmc* resulting in complete clearance within 3 days.

Although the bacterial burden data was acquired from whole eyes and not individual corneas, it is unlikely that *P. aeruginosa* spread to extracorneal sites in the eye, since, by histology, there were no *P. aeruginosa* detected in these areas. Spread of *P. aeruginosa* to the posterior chamber in experimental keratitis has not been described; however, there have been clinical reports of *P. aeruginosa* keratitis evolving into endophthalmitis (22-24).

### The three condensin mutants display unique transcriptional signatures

To gain insight into the origins of the reduced virulence of *P. aeruginosa*, we performed a transcriptomic analysis of condensin deficient cells (GSE161971). We found 1,164 genes, whose expression changed more than 2-fold, compared to the wild type strain, in a statistically significant manner (false discovery rate, FDR, less than 0.1) in at least one of the mutants (Fig. 2A). 534 of these were differentially upregulated and 717 were downregulated (Fig. 2B) with 87 genes being oppositely affected in two of the mutants. Interestingly, most of differentially upregulated genes were unique to each mutant (Fig 2B). Among downregulated genes, a large overlap was observed between *mksB* and *mksB smc* mutants but not with the *smc* strain. Principle component analysis (PCA) further highlighted this relationship revealing a virtually equidistant arrangement of the three strains in PCA coordinates (Fig. 2C). Together, this reveals predominately unique transcriptional signatures for single and double condensin mutants with a distinct overlap between *mksB* and *mksB smc* cells for downregulated genes.

**Figure 2.**
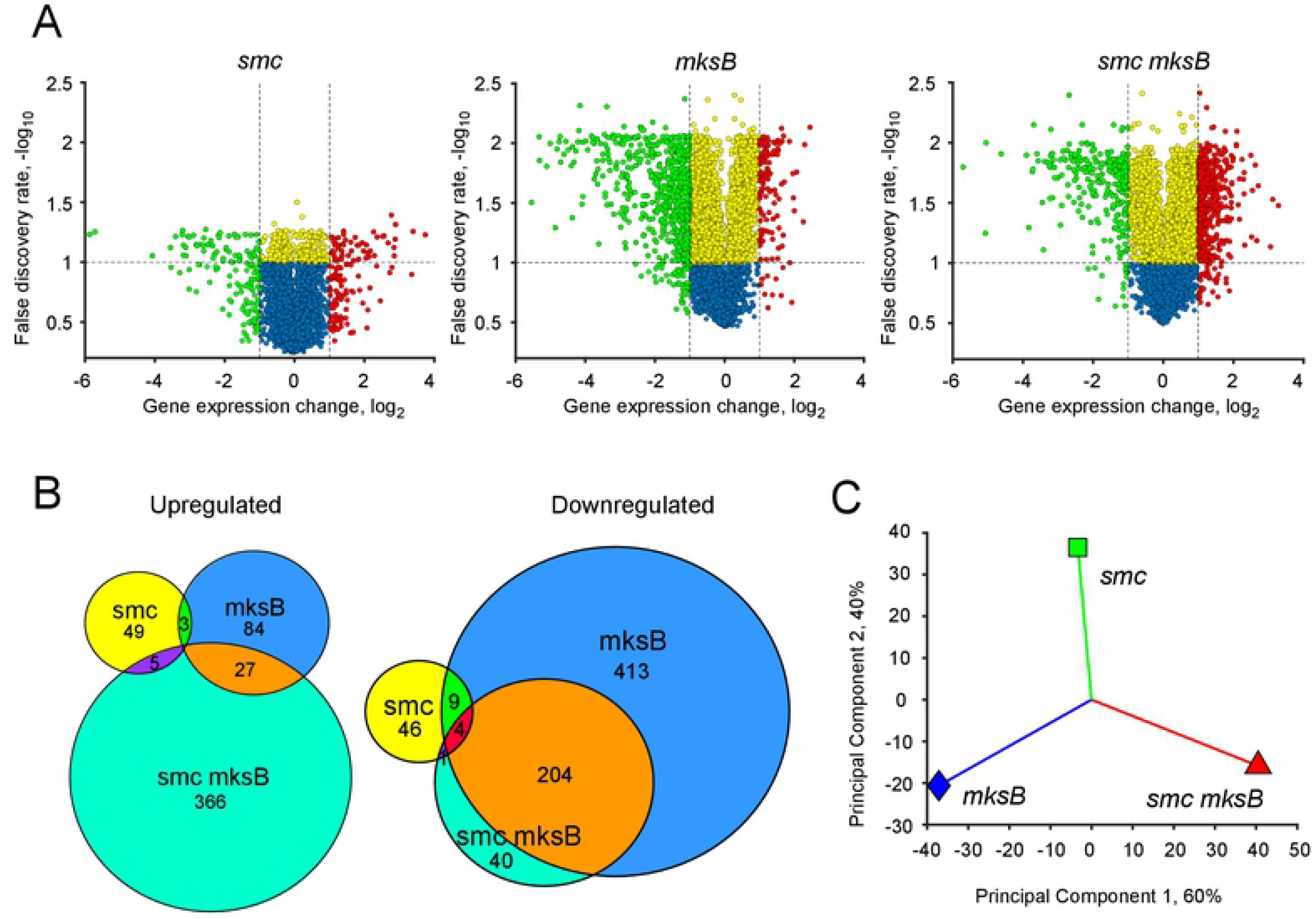
Differential expression of condensin mutants. (**A**) Volcano plots of condensin mutants comparing the expression changes and flase discovery rates for all genes. Downregulated and upregulated (≥ 2 fold) genes are shown in green and red, respectively. Yellow and blue mark the rest of the genes with the FDR values less than or greater than 0.1, as appropriate. (**B**) Venn diagrams of signficiantly (adjusted p-value ≤ 0.01) upregulated (>2 fold; left) or downregulated (≥ 2 fold) genes. (**C**) Principle component analysis of condensin mutants.

### Differentially regulated pathways are dominated by lifestyle and virulence

We next analyzed gene expression using hierarchical clustering. Linkage maps were constructed based on expression changes across all three mutants followed by ranking of the genes according to the separation distance (Fig. S1A). The number of significant clusters in the data set was estimated using the elbow method (Fig S1B). Most of the genes in significantly affected clusters were downregulated in *mksB* and *smc mksB* cells but were mixed in *smc* mutants (Fig. S1A).

Ten out of the 29 statistically identified clusters consisted of artificially separated outliers and were merged with functionally similar groups of genes. Overall, we recognized 18 distinct groups of differentially affected genes (Fig. 3A). One third of the genes (31%) in the clusters were of unknown function or associated with various aspects of primary metabolism (8%; Fig. 3B). For the remaining genes, the response was dominated by lifestyle and virulence pathways.

**Figure 3.**
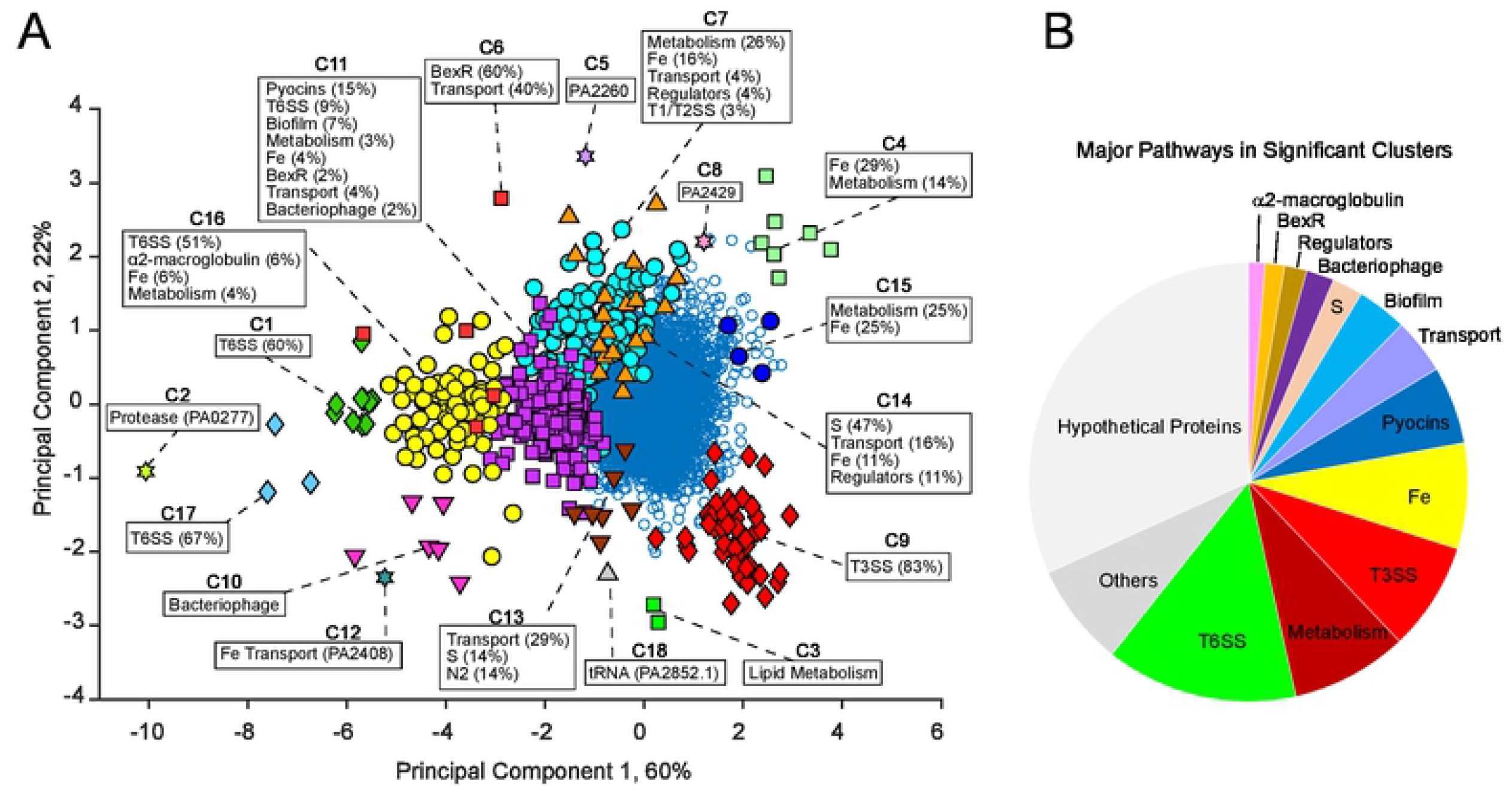
Cluster analysis of expression changes. (**A**) Significant gene clusters identified in condensin mutants. The major pathways comprising each cluster are indicated. (**B**) Pie diagram of pathways identified in significant clusters.

#### Sessile growth

Consistent with the previous physiological analysis (10), genes involved in biofilm formation were prominently present among affected pathways (18 genes, mostly in cluster C11; Table S1). Of these, two *pel* and nine *psl* exo-polysaccharide genes, a major component of biofilms, were identified. Interestingly, a gene encoding hemagglutinin (PA0041; cluster C16) was greatly affected for two condensin mutants. Hemagglutinin is a large filamentous adhesion protein known for its ability to cause red blood cell agglutination and also directly contributes to biofilm formation (25, 26).

#### Iron and sulfur

are critical microelements required for bacterial growth, especially during competitive growth or infection. Besides being key components of the respiratory chain, iron also serves as a signal for biofilm development (27) whereas sulfur is required for cysteine and methionine biosynthesis. *P. aeruginosa* harbors multiple iron acquisition pathways, many of which altered their expression, either upward or downward, in condensin mutants. 38 genes from iron metabolism were found predominantly in clusters C7, C11, and C16. In particular, all pyochelin biosynthesis genes were significantly expressed in cluster 7 (11 genes). Fifteen pyoverdine genes were found significantly expressed across several clusters. Seven of these genes were grouped together in C11 and four in C16. Pyocyanin, a major virulence factor in *P. aeruginosa* which also acts in iron acquisition (28), revealed only 3 significantly affected genes in C4, C7 and C11. Of these, *phzB1* was affected greater than 3.5 fold for all mutants. Nine other iron uptake genes were identified in clusters C7, C15, and C16 including a *tonB2* receptor gene, two ferric iron transporters, *exbB1* and *exbD1*, and *vreI*, a regulator involved in both lifestyle switching and iron uptake (29). Sulfur metabolism genes were abundant and mostly localized in cluster C14 (9 genes) with one gene affected in C11 and one in C13. Tentatively related to respiration were 4 genes from the denitrification pathway (clusters C11 and C7), which contribute, among other functions, to electron transport.

#### Virulence factors

unexpectedly, comprised a distinctive part of the transcriptional response to condensin inactivation. Interestingly, most genes in clusters C16 and C9, found at opposing extremes of the PCA plot (Fig. 3) were assigned to Type 6 and Type 3 secretion systems, T6SS and T3SS, respectively. T3SS genes formed a compact, tightly organized cluster while T6SS genes showed somewhat greater variation. T3SS and T6SS are typically expressed during acute and chronic stages of infection respectively (30) and are essential for *P. aeruginosa* virulence (31, 32).

Besides these two highly specialized systems, several others were affected. The entire operon encoding an α2 macroglobulin homologue (*magABCDEF)* was identified in significant clusters (5 genes in C16 and one gene in C11). These proteins presumably contribute to host evasion through emulation of the human α2-macroglobulin by trapping and inactivating external proteases (33). Cluster 11 also included twenty-eight S, F and R type pyocin genes. Pyocins are bacteriophage-like proteins that display numerous activities, including channel formation, DNA or tRNA degradation and contribute to inter-strain dominance in biofilm communities (34, 35). They can also induce autolysis and thereby enhance biofilm formation (35, 36). Three hydrogen cyanide producing genes, *hcnA, hcnB* and *hcnC* were found in clusters C16 and C11. C16 also contained a hemolysin PA2462, whereas the sole gene in C2 turned out to be a predicted Zn-dependent protease, PA0277 (37).

#### BexR regulon

(7 genes), comprised the core of cluster C6 and part of the larger cluster C11. BexR (PA2432) regulon is expressed in a bistable manner, creating population diversity in a *P. aeruginosa* culture, and encodes several virulence factors such as the toxin AprA, the stress hydrolase PA1202, Lipase A, and the MexGHI-OpmD efflux pump among others (38).

#### Bacteriophage Pf1

Bacteriophage genes were affected including bacteriophage Pf1 implicated in DNA lysis which releases extracellular DNA, a component of biofilm formation (39). Bacteriophage Pf1 genes were found isolated in cluster C10 (6 genes) as well as cluster C11 (4 genes) indicating two distinctly regulated populations.

#### DNA metabolism

Only 2 genes involved in DNA maintenance and repair, *ligD* and PA2150, have been found within significantly affected clusters (C7). These are both related to non-homologous end joining and showed downregulation in both the *smc* and *mksB* mutants. The opposite would be expected were this a part of a DNA damage response.

#### Nutrient metabolism

42 metabolic genes from significant clusters belonged mostly in pathways related to amino acid (11 genes), lipid (5 genes), carbon (11), and nucleotide (3 genes) metabolism. Affected metabolic genes were widely distributed across diverse pathways, suggestive of a dysregulation in gene expression rather than a concerted response. In particular, the 11 genes involved in amino acid metabolism were associated with biosynthesis of arginine, histidine, lysine, tyrosine, valine, leucine and isoleucine.

#### Transporters

39 genes representing 22 ABC (ATP binding cassette) transporters were scattered across several clusters (Table S1). These protein machines are typically dedicated to acquisition or efflux of specific substrates. The affected transporters were associated with diverse functions including import of metals, sulfur and various nutrients, osmoregulation, and export of exopolysaccharides. In addition, three RND transporters, *mexH, mexV*, and *mexW*, associated with pyocyanin and multi-drug efflux (40, 41) changed their expression in at least one mutant greater than 2 fold (FDR < 0.1).

#### Regulators

We identified 28 transcriptional regulators in significant clusters (Table S2) and 56 more were found among genes whose expression changed more than two-fold in a significant manner (FDR less than 0.1) in at least one of the mutants (Worksheet Regulators in Table S3). Most of them belonged to pathways that were significantly affected by the mutations. In particular, 8 and 4 genes, respectively, were involved in the regulation of T3SS and T6SS. BexR and a hemolysin repressor PA2463 (42) were also on this list, whereas four other genes were associated with iron acquisition. Five of the genes, *siaA, siaD, vreI, rsmY* and *qscR* are involved in global control of lifestyle behavior in *P. aeruginosa* including c-di-GMP signaling and biofilm formation pathways. Finally, five putative regulators have not been characterized.

### Transcriptional signature of condensin mutants

Affected regulons were associated with condensin mutations in a distinctive manner (summarized in Fig 4 and Table S3). Only one pathway, bacteriophage Pf1 genes, was similarly affected in all three mutants. The others gave rise to three unique transcriptional profiles.

**Figure 4.**
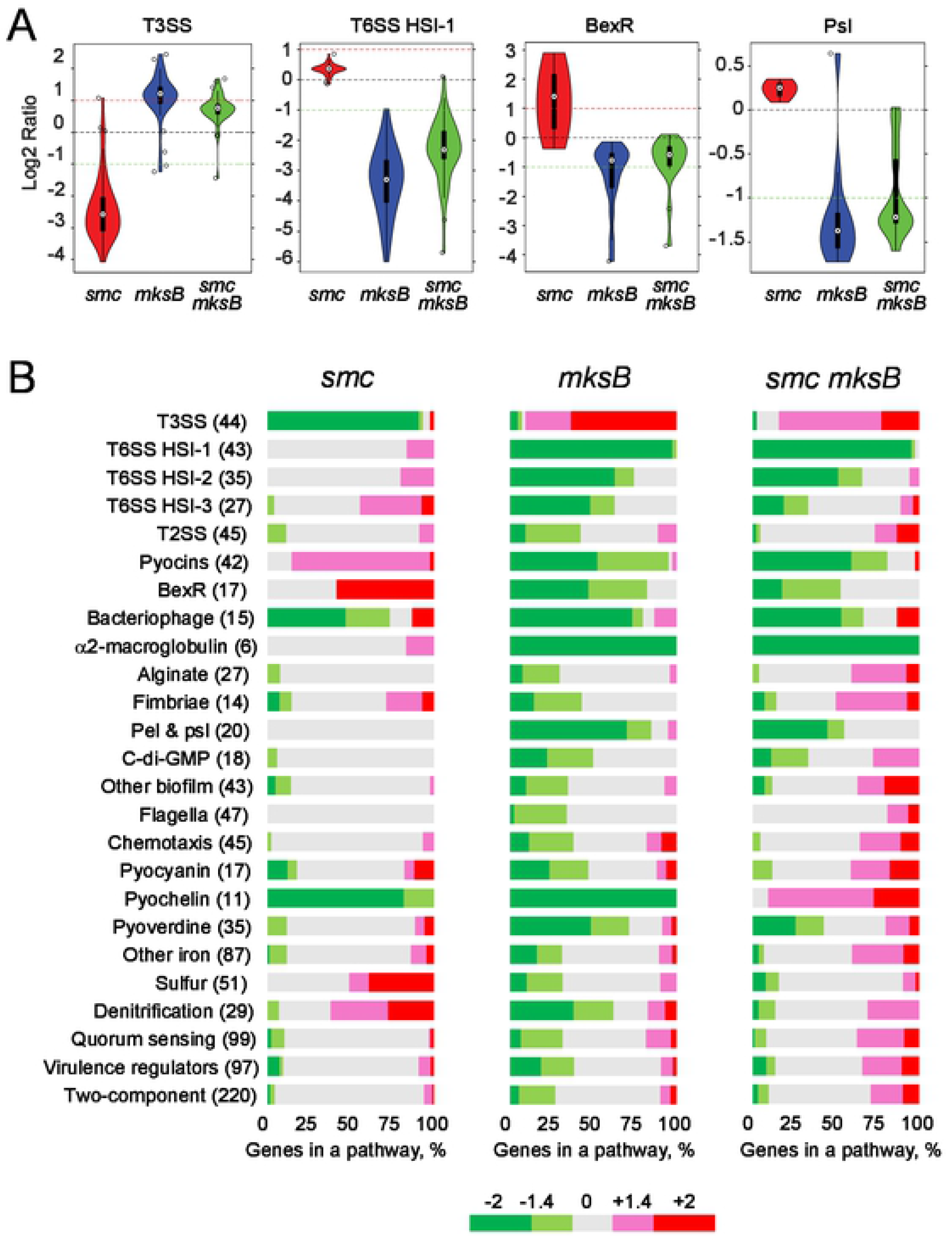
Transcriptional profiles of condensin mutants. (**A**) Violin plots of gene expression changes in select pathways. The black dashed line represents the no change, red line represents ≥ 2 fold upregulation, and green line represents ≥ 2 fold downregulation. (**B**) Summary of affected pathways for each condensin mutant. The number of genes within each pathway is shown in parentheses; genes assigned to each pathway are listed in Tables S3. Bars indicate percent of genes within a given pathway that changed expression at least at the indicated level.

The *smc* strain was characterized by an almost complete downregulation of T3SS accompanied by a broad if mild upregulation of T6SS, BexR regulon, pyocin, hemolysin and α2-macroglobulin production, denitrification, and sulfur assimilation. Iron assimilation was mostly upregulated with a notable exception of pyochelin biosynthesis. Biofilm genes were barely affected except for the mildly induced hemagglutinin (Table S1) and the *psl* regulon, which showed a small but broad upregulation of all genes including the polysaccharide transporter *pslD* (Fig 4A). Taken together, these factors might explain the high surface adhesion of SMC-deficient cells.

The *mksB* regulation pattern was virtually opposite to that of *smc*. As expected for planktonic cells, biofilm related genes were downregulated, as were iron and sulfur uptake and denitrification. Among virulence factors, T3SS was induced, while substantial downregulation was observed for BexR, T6SS, pyocins, bacteriophage Pf1, hemolysin, hemagglutinin, α2-macroglobulin, fimbriae, the PA0277 protease, and hydrogen cyanide toxin genes (Table S1).

The *smc mksB* cells displayed similar regulation patterns to *mksB* for many growth and secretion pathways. These included upregulation of T3SS and substantial downregulation for T6SS and biofilm genes. Significant downregulation was observed for BexR, bacteriophage, α2-macroglobulin, PA0277 protease and hydrogen cyanide producing genes. However, the regulation patterns were more ambiguous than for *mksB* cells, with many regulons containing multiple up- and downregulated genes. Moreover, certain cases could not be explained by a combination of SMC- and MksB-dependent regulation. For example, pyochelin production was inhibited in both single condensin mutants but upregulated in the double mutant strain.

### Expression of genes and their regulators partially correlates

Inactivation of condensins altered the expression of numerous transcription factors and signal transduction pathways. This raised the possibility that condensins could exert their transcriptional control through specific transcriptional regulators. To explore this possibility, we compared expression changes of affected regulons and their immediate regulators. We indeed observed a good correlation between the two, but only in a subset of cases (Fig. 5).

**Figure 5.**
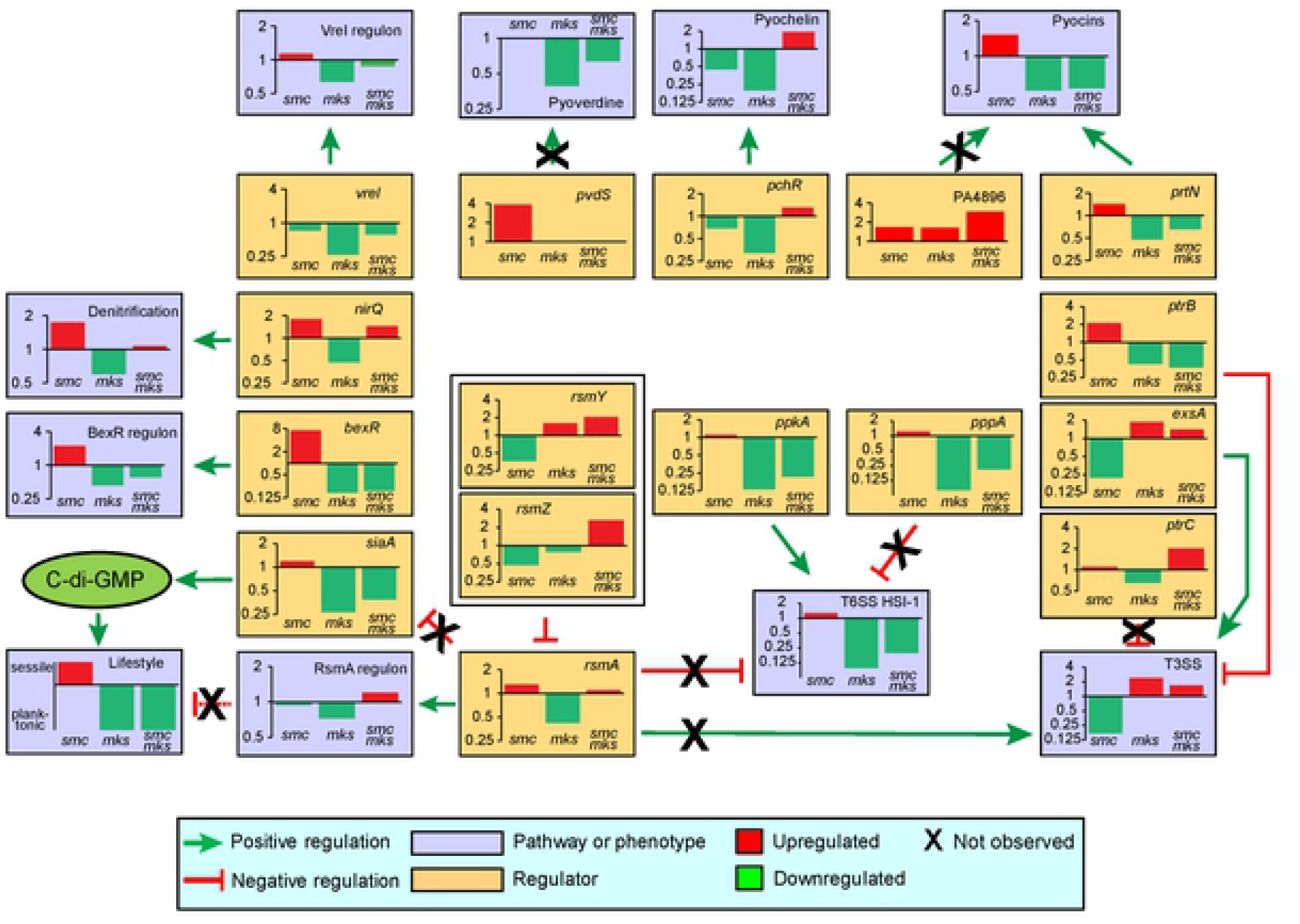
Comparison of expression changes of affected regulons (slate boxes) and their primary regulators (tan boxes). Expression changes for each mutant are shown as bar graphs in each box. Arrows indicate the known regulatory connections between the genes. Crosses mark regulatory pathways that are inconsistent with the gene expression changes found in condensin mutants.

A strong correlation was observed for pyochelin biosynthesis genes and their regulator PcrR (43), BexR and its regulon (38), VreI and its regulon (29), and denitrification genes controlled by NirQ (44). In all three mutants, the average expression changes of the entire regulon paralleled those of the cognate regulator (Fig. 5). In contrast, no correlation for any of the mutants was found for the pyoverdine production genes and *pvdS*. For pyocin production, significant expression changes were observed for two of its regulators, PrtN and PA4896 (29, 45). However, only one of them, PrtN, matched the behavior of the regulon. Likewise, the expression pattern of T6SS matched that of its activator PpkA (46) but not of the antagonists PppA (46) and RsmA (47) whereas the T3SS expression mirrored the levels of the activator ExsA (48) and repressor PrtN (45) but not PtrC (49) or RsmA (47, 48).

Two pathways could explain the sessile/planktonic behavior of the condensin mutants. The switch between these two lifestyles is often controlled by the c-di-GMP signaling network (50). c-di-GMP biosynthesis occurs through the siaA/D protein system which is in turn, negatively regulated by free RsmA, that is the fraction of RsmA that is not associated with its inhibitor RNAs, *rsmY* and *rsmZ* (47). We found large changes in the expression of SiaA, which correlated with the dominance of the sessile and planktonic phenotypes observed in condensin mutants (Fig. 5). In contrast, the expression of the RsmA regulon, which includes a number of lifestyle genes, showed little correlation with the propensity of condensin mutants for biofilm formation.

In summary, expression changes in the affected regulons and their immediate regulators often paralleled each other, suggesting that at least some of the changes were not caused by a direct transcriptional control from condensins but rather propagated through the signaling network of *P. aeruginosa*.

### SMC deficiency impairs amino acid biosynthesis and resistance to peroxide

Transcriptomic analysis revealed many physiological deficiencies that could explain the reduced pathogenicity of condensin mutants. We first examined the mutants for fitness defects. We did not find notable growth abnormalities in a rich medium (Fig. 6A). In M9 medium, however, growth of *smc* but not *mksB* or *smc mksB* cells was markedly delayed (Fig. 6B). This lag varied in magnitude but remained prominent when trace elements (Fig. 6C) or amino acids (Fig. 6D) were added into the medium, suggesting that it is caused by multiple factors. The lag was indeed caused by an absence of SMC since it was largely abolished by an extrachromosomal expression of the gene (Fig. 6E).

**Figure 6.**
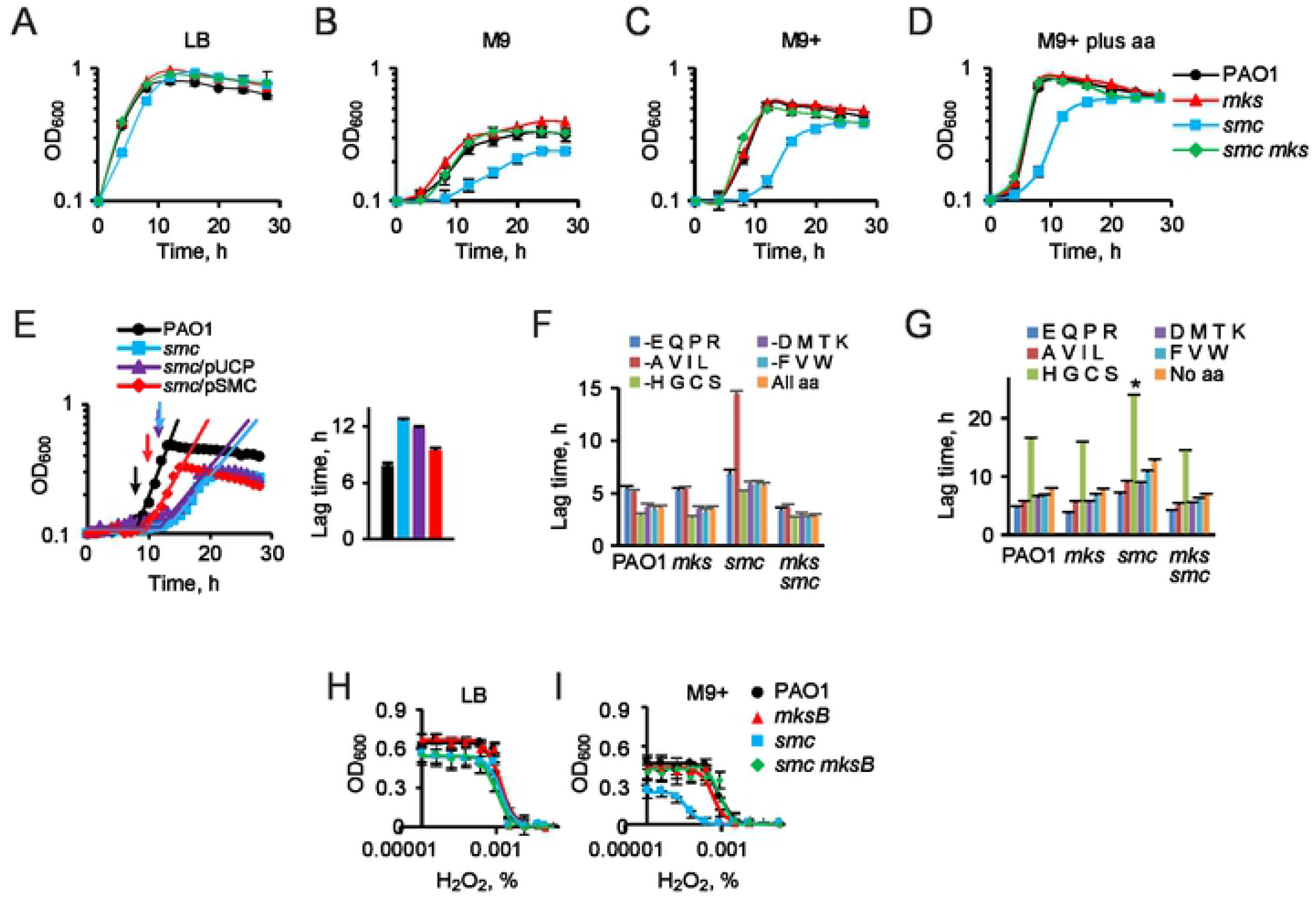
Fitness of condensin variant cells under nutrient limiting conditions. (**A**-**D**) Growth of *P. aeruginosa* and condensin mutants in LB (**A**), M9 medium (**B**), M9 medium supplemented with trace ions (**C**) and with all amino acids (**D**). Cells were inoculated at OD_600_ of 0.1. (**E**) Complementation analysis of *smc* phenotype. Growth curves of cells that lack the *smc* gene or express it from the chromosome (PAO1) or a plasmid (pSMC). The growth curves (±SD, N=3) were fit to Eq. 1 (straight lines), and the best-fit lag times are shown on the graph on the right. Arrows indicate the end of the lag. (**F, G**) The best-fit lag times for cells grown in M9+ medium lacking only the indicated amino acids (**F**) or supplemented with them (**G**). Asterisk marks conditions where no bacterial growth was observed. (**H, I**) Hydrogen peroxide susceptibility of condensin variant strains in the indicated growth medium (±SD, N = 3).

To determine which amino acids contributed to the lag, we grouped them into five sets according to their biosynthetic mechanisms, and omitted one of each group from the growth medium. To facilitate the comparison across multiple conditions, we measured the lag and growth rate for each condition as illustrated in Figure 6E. The omission of any of the amino acid groups delayed the growth of *Δsmc* cells; however, the absence of hydrophobic amino acids (alanine, valine, isoleucine and leucine) had a particularly strong effect (Fig. 6F). In contrast, the growth rate of the *Δsmc* cells, but not the other mutants, was equally reduced in the absence of any set of amino acids (Fig. S1A). The addition of individual sets of amino acids into the minimal medium reduced the lag of all tested bacteria but did not eliminate the distinctive delay in growth of *Δsmc* cells (Fig. 6G, S1B). The only exception observed was for the mixture of glycine, cysteine, serine and histidine, which increased the lag instead of decreasing it. The increase was due to cysteine (Fig. S1C, D), presumably caused by its iron chelating activity (51). Even in this respect, the fitness of SMC-deficient cells was notably reduced.

We did not find any increase in susceptibility of condensin deficient *P. aeruginosa* to lysozyme or defensins β-defensin and lactoferrin. However, *Δsmc* but not *ΔmksB* or *ΔsmcΔmksB* cells were susceptible to hydrogen peroxide, a mimic of phagocyte-produced nitric oxide (Fig 6H, I). The increase in susceptibility was clear in minimal medium (Fig. 6I) but not LB (Fig. 6H), suggesting that it might have common roots with the decreased fitness of condensin deficient cells.

### Inactivation of MksB and SMC reduces production of virulence factors

The main pigments of *P. aeruginosa*, pyocyanin and pyoverdine, are among the more abundant of its virulence factors and frequently contribute to initial host colonization by *P. aeruginosa* (52-55). We found that production of the two pigments mirrors the other phenotypes of condensin mutants (Fig. 7A). Namely, production of pyocyanin was only marginally increased, if at all, upon inactivation of SMC but was virtually undetectable in *mksB* and *smc mksB* cells (Fig 7B). In contrast, pyoverdine production was not significantly affected in any of the condensin mutants (Fig 7C). This phenotype was found to be indeed associated with *mksB* after deleting the gene using a degron system (Fig. 7D). To this end, the endogenous *mksB* was replaced with its DAS4-tagged version in a strain that lacks a ClpXP adaptor protein SspB. SspB was then expressed from an externally delivered plasmid. Degradation of MksB-DAS4 using the degron system abolished pyocyanin production by *P. aeruginosa* (Fig. 7E).

**Figure 7.**
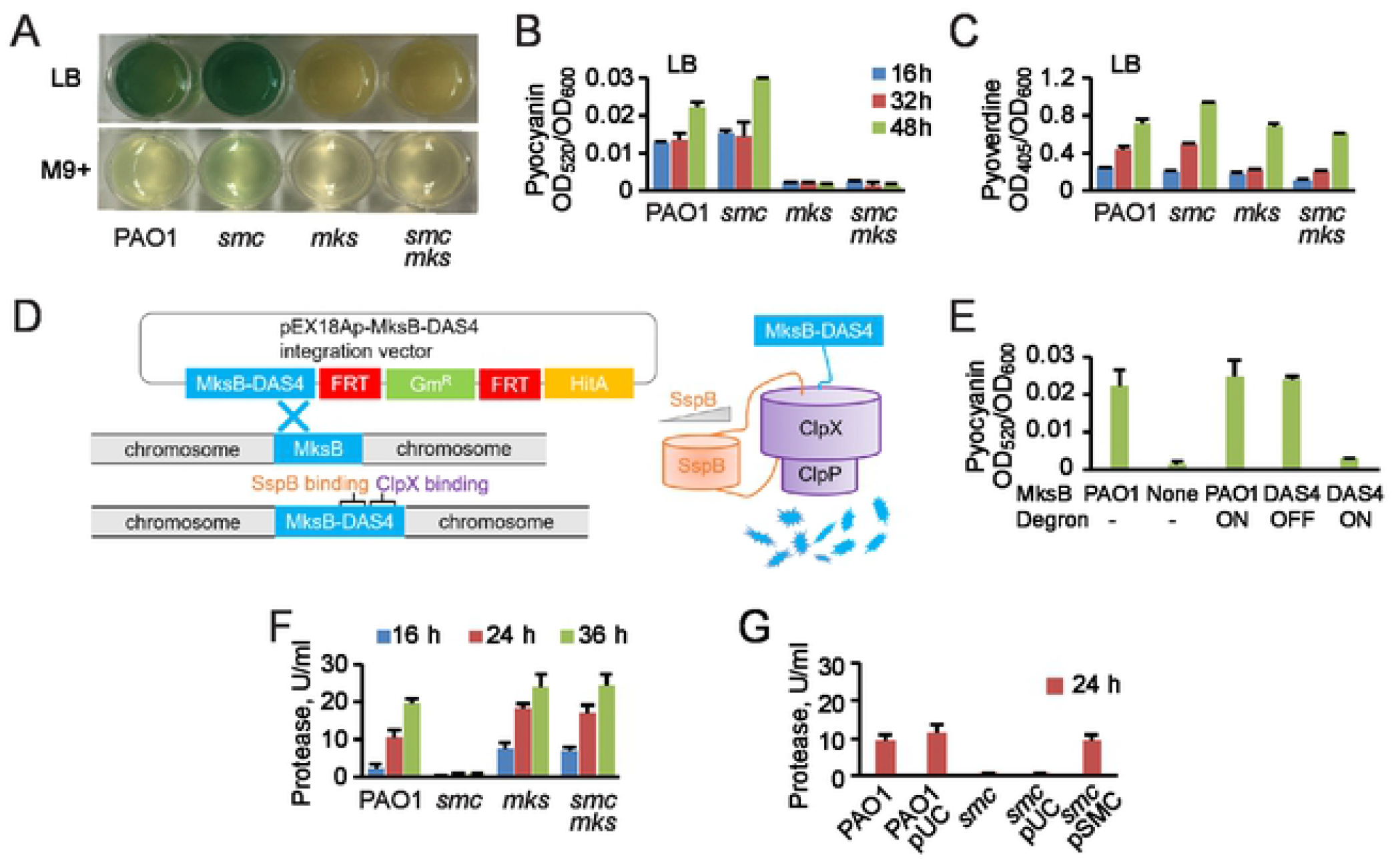
Virulence factor deficiency of *P. aeruginosa* PAO1 and condensin mutants. (**A**) Bacteria grown at 37 °C in LB for 2 days or M9 medium supplemented with trace ions (M9+) for 3 days. (**B, C**) Production of pyocyanine and pyoverdine, as indicated (±SD, *n* = 2), by cells grown in LB. (**D**) A diagram illustrating depletion of MksB using the degron system. (**E**) Inducible depletion of MksB impairs pyocyanin production. Pyocyanin (±SD, *n* = 2) was measured for PAO1 (WT), *mksB* (None), or OP132 (Δ*sspB mksB*::*mksB-DAS4*; DAS4) cells that were transformed with the pUCP22 (OFF) or pSspB (ON) plasmid. (**F**) Extracellular protease production (±SD, *n* = 3). (**G**) Complementation analysis of the secreted protease activity in PAO1 and *smc* cells that harbor, when indicated, plasmids pUCP22 or pUCP22-SMC (±SD, *n* = 3).

Unlike with the pigments, protease secretion was reduced in SMC-but not MksB-deficient cells (Fig. 7F). This deficiency was completely restored by extrachromosomal production of SMC (Fig. 7G). This result mirrors the downregulation of T3SS observed for *smc* mutants (Fig.4).

### Condensins are integrated into P. aeruginosa *regulatory network*

We next determined whether condensins act independently or as part of a regulatory network. To this end, we explored genetic interactions between condensins and three regulators, *rsmZ*, VreI and RhlI. The first two regulators were strongly affected by condensin deletions (Fig. 5), whereas the third one showed only borderline changes. All three regulators were implicated in the control of planktonic and sessile behavior, each via its own mechanism (29, 56, 57). As a proxy for these phenotypes, we followed competitive growth of the mutants in the presence of their parental PAO1 strain as well as their propensity to stick to a surface.

Under our assay conditions, cells devoid of the quorum sensing regulator RhlI were indistinguishable from the parental strain (Fig. 8A, B). The deletion of *rsmZ*, a negative regulator of RsmA, had no effect on the surface adhesion phenotype of any of the condensin variants (Fig. 8B) but modestly impaired their competitive growth (Fig. 8A). No clear evidence of genetic interaction between *rsmZ* and condensins was detected. Curiously, however, an interaction was found for VreI, a transcriptional factor that is primarily involved in iron acquisition (29). The deletion of *vreI* increased the biofilm producing ability of wild type cells to the levels observed for SMC deficient cells but had no effect on the *smc* mutants (Fig. 8B). Not dissimilarly, the inactivation of VreI did not change the competitive growth of the wild type cells but diminished the defects caused by the *smc* deletion (Fig. 8A).

**Figure 8.**
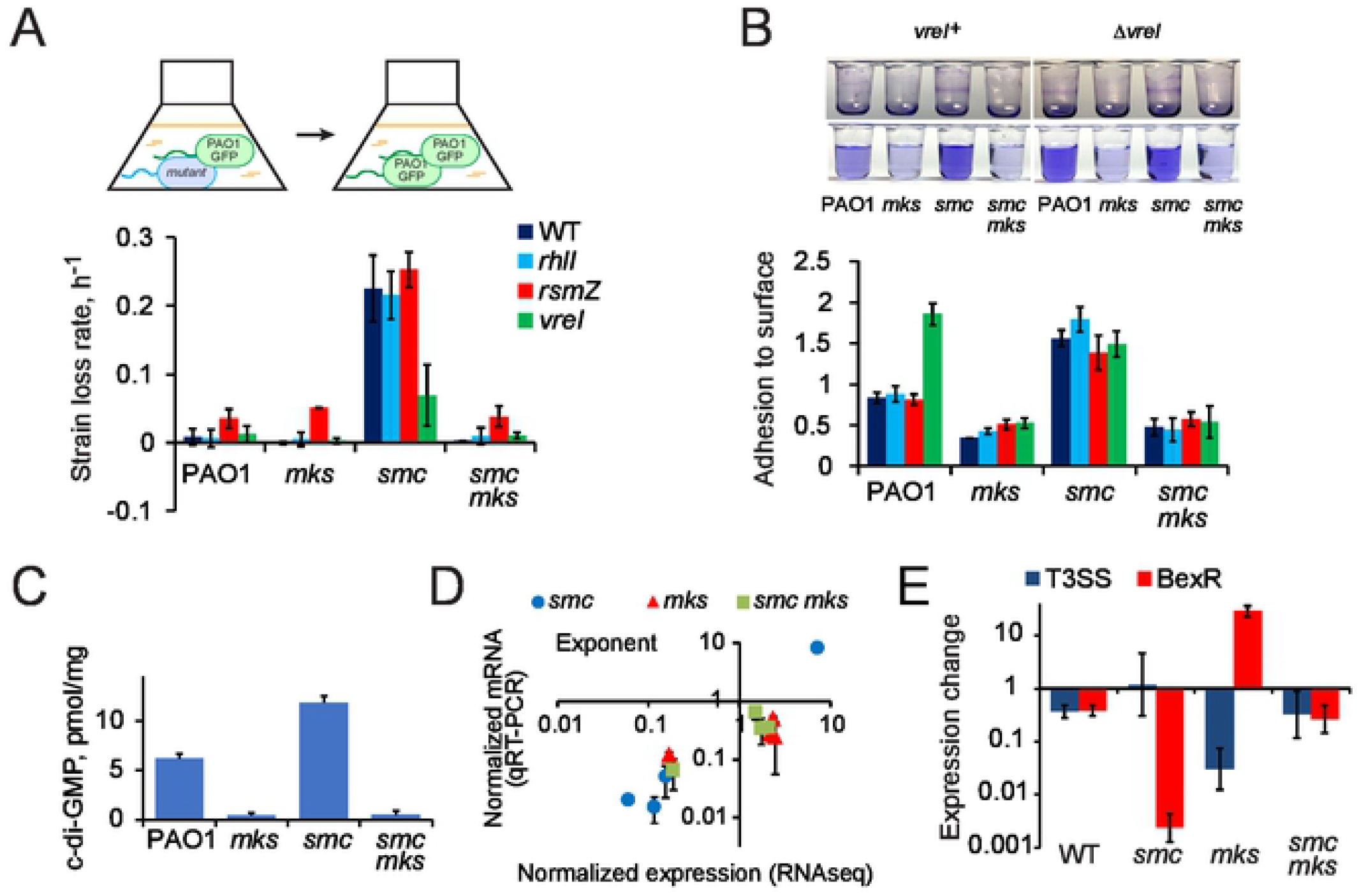
Condensins are integrated into signaling network. (**A**). Strain loss rate during competitive growth of the indicated double mutants of condensins and select virulence regulators and the parental WT strain (± SD; *n* = 3). (**B**). Biofilm formation on PVC by cells deficient in condensins and the indicated virulence regulators (± SD; *n* = 3). (**C**) c-di-GMP levels normalized to the total nucleotide content in condensin mutants. (**D**) Comparison of gene expression changes in the exponential phase cells, normalized to wild type cells, for *spcS, exoT* and *pcrV* and *bexR*, as determined by RNA-seq and qRT-PCR. (**E**) Expression levels of T3SS (average for *spcS, exoT* and *pcrV*) and BexR in stationary phase cells normalized to those in the exponentially growing cells.

Given the starkly different functions of condensins and the VreI regulon, the genetic interaction found between the two proteins suggests that they affect the lifestyle of the bacterium indirectly, by exploiting the existing signaling network. To evaluate this notion, we measured the levels of c-di-GMP in condensin variant cells, which is a secondary messenger responsible for lifestyle switching in *P. aeruginosa* (50). We indeed found that the level of c-di-GMP varied in a condensin-dependent manner. The amount of c-di-GMP was elevated in SMC deficient cells but was virtually undetected in *mksB* and *smc mksB* mutants (Fig. 8C). These intracellular messenger levels are fully consistent with the observed phenotypes suggesting that condensin mutations induce lifestyle switching by triggering the intracellular signaling network.

To further evaluate this conclusion, we examined the effects of condensins on quorum sensing regulation. To this end, we compared condensin-induced gene expression changes in exponential and stationary cells for the BexR regulon and T3SS, which have been implicated in the transcriptomic analysis (Fig. 4A). Should condensins be directly involved in the control of these genes, we expected similar effects on their expression in growing and stationary cells. Using qRT-PCR, we measured the gene expression levels for *spcS, exoT* and *pcrV* from T3SS and *bexR*. For exponentially growing cells, the mRNA levels of these genes from qRT-PCR and RNA-seq measurements were in a good agreement with each other (Fig. 8D). Upon transition to stationary phase, the transcript abundancies changed in a condensin dependent manner (Fig. 8E). The expression of both regulons modestly declined in wild type cells. In SMC-deficient cells, the expression of T3SS remained at the same level as in the exponential phase whereas BexR expression levels substantially declined. In *mksB* mutants, the expression of T3SS declined by almost 40-fold whereas BexR was overproduced by an order of magnitude. The expression pattern in the double mutant was similar to that found in the parental cells. Thus, the regulatory effects of condensin deletions are not additive with the regulation that accompanies the onset of stationary phase.

## Discussion

Condensins are an emerging target for antibacterial drug discovery (58). *P. aeruginosa* condensins were found essential for virulence in a mouse model of lung infection (10). Here, we show that they are also essential during ocular infections. Inactivation of any of the two PAO1 condensins accelerated clearance of *P. aeruginosa* from the infected animals. Mice infected with *smc* cells initially showed a higher bacterial load than those with the parental strain, but that was significantly reduced by Day 2 and continued to decline. The higher initial pathogenicity was likely caused by an upregulation of T6SS, hemagglutinin and exopolysaccharide production in these cells resulting in a higher propensity for biofilm formation (10). The double condensin mutants were the least virulent and were completely cleared by Day 3. In contrast, the double mutant cleared somewhat slower than the SMC-deficient strain during the acute stage of lung infection (10), highlighting a variation in disease models.

Condensins primarily function in global chromosome folding and segregation. *P. aeruginosa* harbors two condensins, which play distinct roles in chromosome maintenance (10, 17). However, deletion of condensins induces only mild structural defects in *P. aeruginosa*, which are unlikely to cause the observed reduction in the virulence of the bacterium.

Besides structural chromosome maintenance, condensins have been recently implicated in lifestyle control (10). SMC deficient *P. aeruginosa* becomes highly prone to biofilm formation and, conversely, poorly performs during competitive growth with its parental strain. In contrast, *mksB* mutants appeared better adapted for planktonic growth. We show here that the regulatory functions of condensin are much broader than lifestyle control and include numerous metabolic and virulence programs. In this respect, condensins can be qualified as global regulators of gene expression.

Transcriptional programs imposed by condensins were largely dissimilar. A substantial overlap was only found for genes downregulated in *mksB* and *smc mksB* cells, whereas upregulated gene sets were unique to each of the three mutants (Fig. 2B). In all cases, the transcriptional response was dominated by lifestyle and virulence programs. MksB deficient cells downregulated multiple systems that promote surface adhesion, making them fit for planktonic growth, while upregulating T3SS (Fig. 4). This regulatory pattern is reminiscent of cells during an acute phase of infection (59, 60). Conversely, *smc* mutants induced several surface adhesion mechanisms, downregulated T3SS and upregulated a variety of host-directed virulence mechanisms including T6SS, pyocins, hemolysin, hemagglutinin and α2-macroglobulin. The induction of multiple iron and sulfur acquisition pathways would also benefit the bacterium in its competition with host cells (46, 47). This regulatory pattern conspicuously resembles the behavior of *P. aeruginosa* during chronic stages of infections (32, 61). Thus, the deletion of *smc* and *mksB* triggers opposite pathogenic programs in *P. aeruginosa*.

Many changes could not be attributed to a lifestyle switch. In particular, the altered expression of metabolic genes (Fig. 3) resulted in amino acid dependency and hydrogen peroxide susceptibility of *smc* mutants (Fig. 6). The presence of amino acids in cystic fibrosis sputum was shown to increase the ability of *P. aeruginosa* to establish the disease (62), indicating that such metabolic deficiencies are not conducive to pathogenesis. In this respect, transcriptional changes caused by the deletions of condensins bear elements of dysregulation. The observed interference between condensin mutations and the normal signal transduction (Fig. 8) further corroborates this conclusion.

Other DNA packing proteins have been previously shown to affect gene expression. For example, HNS was implicated in silencing horizontally acquired genes (63). The case of condensins is different because of its modest copy number, which is approximately 100 molecules of SMC and 3,000 molecules of MksB per cell (10). It seems unlikely, therefore, that condensins affect gene expression through a direct binding at the affected promoters. Rather, their absence could be altering the overall chromosome structure with downstream global effects on transcription. Alternatively or in parallel, SMC and MksB could be directly involved in regulation of key members of the regulatory virulence network. The high correlation between the expression of many pathways and their immediate regulators (Fig. 5) is consistent with the latter mechanism.

The implications of this result are broad. At a minimum, these data reveal that the lifestyle decisions in *P. aeruginosa* are dependent on the activity of its condensins and by extension the global structure of its chromosome. Exploiting these interactions will likely yield new antibacterial targets and therapeutic strategies. Looking broader, these findings implicate structural chromosome dynamics in the adaptive and pathogenic behavior of bacteria and the operation of its regulatory virulence network.

## Methods

### Ethics Statement

*Mus musculus* were used in these experiments. All procedures were conducted according to the guidelines and recommendations of the Guide for the Care and Use of Laboratory Animals, the University of Oklahoma Health Sciences Center Institutional Animal Care and Use Committee and Office of Animal Welfare Assurance, and the Association for Research in Vision and Ophthalmology Statement for the Use of Animals in Ophthalmic and Vision Research. These studies were approved under protocols 13-054 and 14-084. Male C57BL/6J mice were purchased from a commercial vendor (Stock No. 000664, Jackson Labs, Bar Harbor ME). All mice were acclimated to housing under biosafety level 2 conditions for at least one week to equilibrate their microflora. Mice were used in experiments at 6-8 weeks of age.

### Murine *Pseudomonas aeruginosa* Keratitis

Prior to infection, each mouse was anesthetized with a combination of ketamine (85 mg/kg body weight; Ketathesia, Henry Schein Animal Health, Dublin, OH) and xylazine (14 mg/kg body weight; AnaSed; Akorn Inc., Decatur, IL). Experimental *P. aeruginosa* keratitis was induced in mouse eyes by scratching the corneal epithelium with an 18-gauge needle and immediately topically applying 5 µl of brain heart infusion media containing approximately 10^6^ CFU of *P. aeruginosa* PAO1, the MksB-deficient mutant (MksB-), the SMC-deficient mutant (SMC-) or the double mutant deficient in both condensins (MksB-/SMC-). The inocula were allowed to incubate on open eyes for 5 min. The contralateral eye served as the uninjected control. Inocula were quantified by track dilution as noted below.

Ocular growth of *P. aeruginosa* in whole eyes of C57BL/6J mice was quantified as follows. Whole eyes were removed from euthanized mice at 1, 2, and 3 days postinfection. Eyes were homogenized in PBS with sterile 1mm glass beads (BioSpec Products, Inc., Bartlesville OK). Homogenates were plated and bacterial numbers were estimated by the track dilution method (64, 65). Values represent the mean ± SEM for N≥5 eyes per time point. At least two independent experiments were performed. The limit of detection for this assay was 10 CFU.

To analyze gross corneal pathology, mice were anesthetized with isoflurane and eyes were imaged using an operating microscope (Zeiss OPMI Lumera and Medilive MindStream; Zeiss, Dublin CA). Images are representative of at least three eyes per time point from at least two independent experiments. To analyze corneal pathology at the tissue level, eyes were harvested from mice at 1 day postinfection. Harvested eyes were incubated in buffered zinc formalin or Davison’s fixative for 24 h at room temperature (64, 65). Eyes were then transferred to 70% ethanol, embedded in paraffin, sectioned, and stained with hematoxylin and eosin. Images are representative of at least two eyes per time point from at least two independent experiments. All images were analyzed and pathological changes documented by a masked observer.

### Statistics

Data represented are the arithmetic means ± the standard errors of the mean (SEM) of all samples in the same experimental group in replicate experiments, unless otherwise specified. A value of p<0.05 was considered to be statistically significant. The Mann Whitney Rank Sum test was used to compare the unpaired experimental groups to determine the statistical significance for all assays (64, 66). All statistical analyses were performed using Prism 6.05 (GraphPad Software, Inc., La Jolla CA).

### Strains and growth media

Condensin deficient strains *ΔmksB* (OP109), *Δsmc* (OP107) and *ΔmksBΔsmc* (OP115), and their parental strain PAO1 (ATCC47085) were previously described (10). Cells were grown with vigorous agitation (200–220 rpm) at 37°C. M9 minimum medium was supplemented with 0.25% citric as the carbon source. The artificial tear medium (ATM) contained the preservative-free Equate Restore Plus lubricant eye drops supplemented with 33.7 mM Na_2_HPO_4_, 22 mM KH_2_PO_4_, 8.55 mM NaCl, 9.35 mM NH_4_Cl, 0.3 CaCl_2_, 1 mM MgCl_2_, 2 mg/ml lysozyme, 20% human serum (H4522 Sigma-Aldrich), and 0.5% of carboxymethyl cellulose. Unless indicated otherwise, all growth media were supplemented with a mixture of trace ions (Table S4; (67)).

Deletion mutants (Table 1) were constructed using allele replacement. Approximately 500 bp of chromosomal fragments flanking *rhlI* (PA3476), *rsmZ* (PA3621.1) and *vreI* (PA0675) genes were generated by PCR and then spliced together as appropriate using overlap PCR with a linker 5-ATGGCGGCCGCTTAA to generate complete deletions of the genes. The resulting PCR products were inserted between the XhoI and KpnI sites of pEXG2 (68) to generate plasmids pEXG2-ΔrhlI, pEXG2-ΔrsmZ and pEXG2-ΔvreI. These plasmids were conjugationally transferred into recipient *P. aeruginosa* strains to create the desired in-frame deletions (17).

**Table 1.**
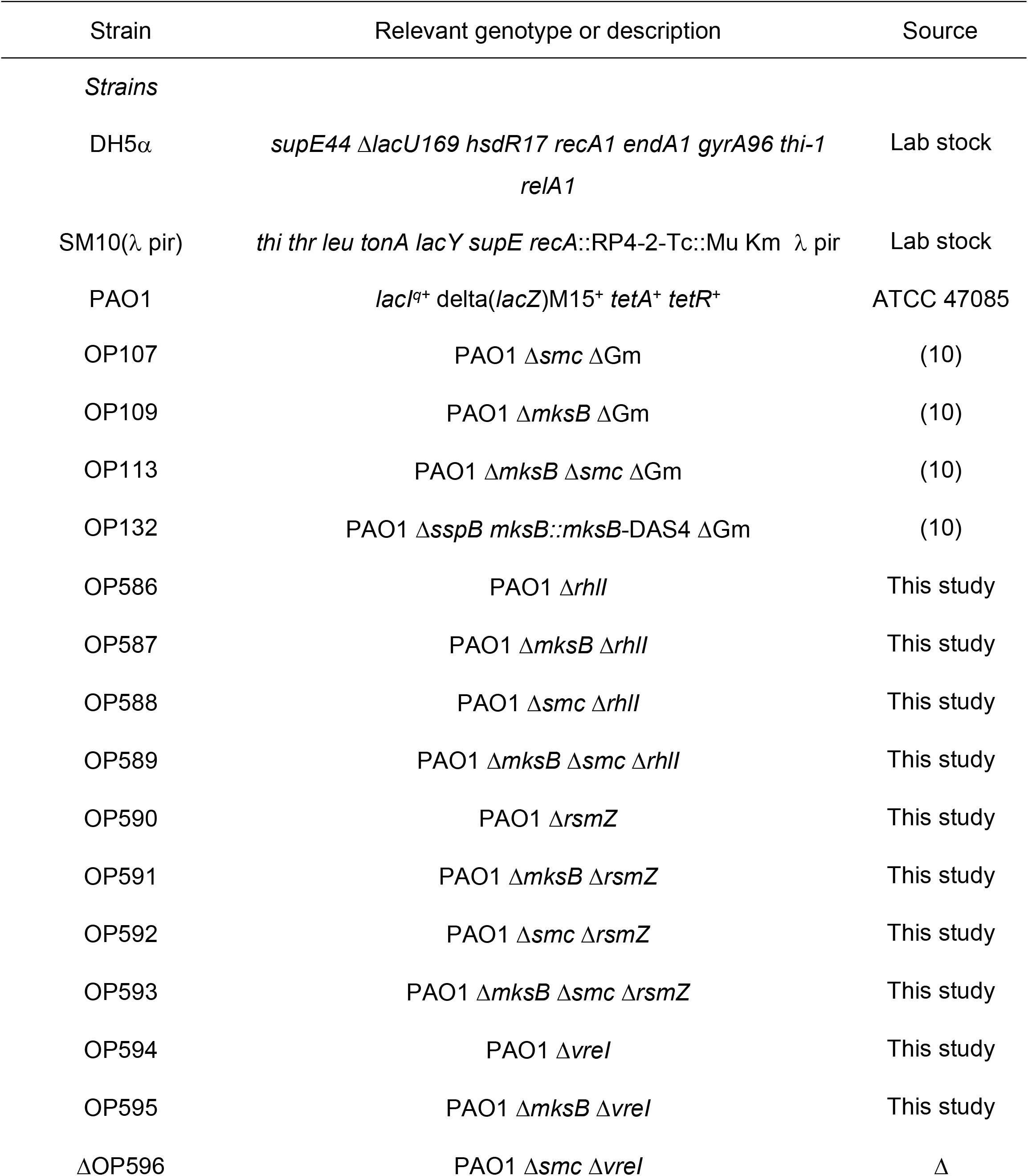
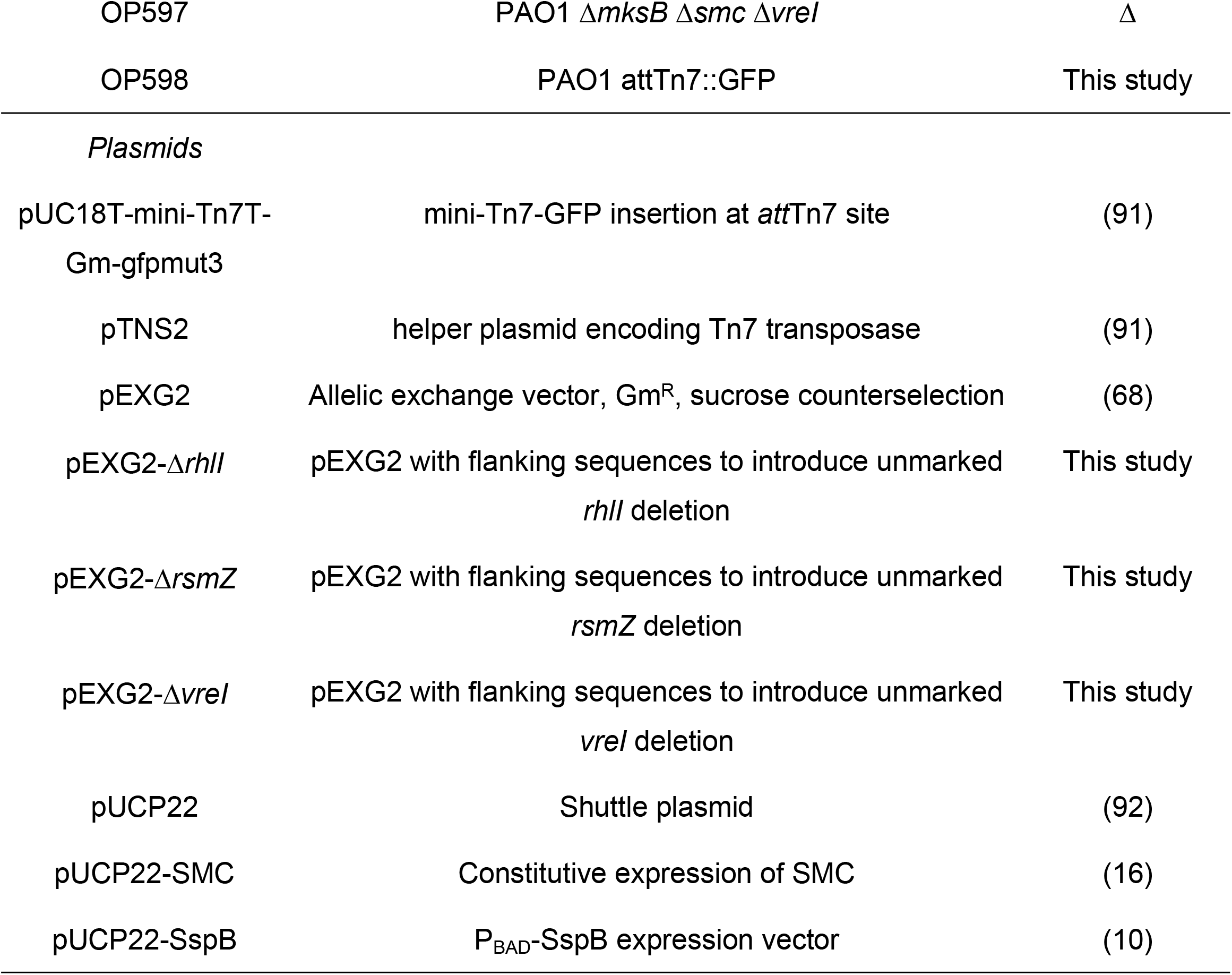
Strains and plasmids

### Microbiology techniques

Overnight cultures were inoculated into LB, grown with aeration to an OD_600_ of 0.6, washed with appropriate minimal medium and then inoculated into microplates containing M9, ATM or LB supplemented with 100 μg/ml of the indicated amino acids at an OD_600_ of 0.1. Amino acids were combined in the following groups: Group 1 (Phe, Tyr and Trp), Group 2 (Asp, Met, Thr and Lys), Group 3 (His, Gly, Cys and Ser), Group 4 (Ala, Val, Ile and Leu) and Group 5 (Glu, Gln, Pro and Arg). The cell growth was followed using a Tecan Spark 10M microplate reader. The lag time, *t*_*lag*_, and growth rate, *r*, were determined by fitting the growth curves to the equation

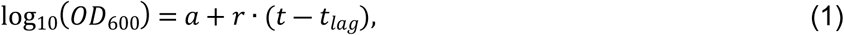

where *a* is the logarithm of the initial optical density of the culture. The doubling time was then determined for the best-fit growth rate as *t*_*1/2*_ = log_2_(10)/*r*.

Minimal inhibitory concentrations (MIC) were measured using the two-fold serial dilution method. 10^4^ exponential phase cells in 100 μl were dispensed into microplates and incubated at 37oC for up to 24 hours with shaking at 180 rpm.

Production of pyocyanin and pyoverdine was quantified as previously described (69, 70). Cells were grown overnight in LB with shaking at 37 °C, diluted 100-fold into fresh LB medium in a 24-well plate, and grown as a standing culture at 37 °C for the indicated time. The cells were then pelleted, and the concentration of pyoverdine in the supernatant determined from its absorbance at 405 nm (70). Concentration of pyocyanin was determined following chloroform-HCl extraction of the supernatant based on its absorbance at 520 nm (69).

### Intracellular c-di-GMP concentration

The c-di-GMP concentrations in PAO1 and condensins mutant strains were measured using the Cyclic-di-GMP Assay Kit (LUCERNA, Catalog Number: 200-100) with modification. Briefly, overnight cells were diluted 1,000-fold in LB and grown for 16 hours at 37°C. 3 ml cultures were then pelleted, resuspended in 300 μL of ice-cold extraction buffer (acetonitrile/methanol/ DEPC-treated water, 4:4:2, v/v/v), and the total nucleic acid extracted as previously described (71) and resuspended in 300 μL of DEPC-treated water. 50 µl of the samples were supplemented with a serially diluted c-di-GMP standard and the assay reactions set up according to the manufacturer’s instructions. After 30 min incubation, the rate of fluorescence increase at 482/505 nm was measured, and the concentration dependence of the rates used to calculate the concentration of c-di-GMP in the unsupplemented samples. The concentrations were then normalized to the total protein content in the samples, which were measured using Bradford assay following the suspension of 500 μl cell culture in 400 μL of 0.1 M NaOH and heating it for 15 min at 95 °C.

### Protease secretion

Cells were grown in LB overnight at 37°C. When appropriate, the cells were pelleted and 150 µl supernatant was added to 350 µl freshly prepared reaction buffer (0.1 M Tris-HCl pH 8.0, 1% azocasein and 0.5% NaHCO_3_). The mixtures were incubated for 20 minutes with shaking at 37°C and supplemented with 1 ml 7% ice-cold perchloric acid. After a brief centrifugation, the cleared supernatant from the solution was added to 150 µl of 10 M NaOH and the OD at 430 nm was measured. One protease unit was defined as the amount of the enzyme that yields an increase of 1 OD unit per hour.

### RNA isolation

Cells were grown in LB at 37°C until late exponential (OD_600_ of 0.6) or stationary (OD_600_ of 3) phase, as indicated. RNA was isolated using the hot phenol method as previously described (72). The extracted RNA was resuspended in DEPC-treated water at 0.2 mg/ml, adjusted to 1xDNase I buffer (10 mM Tris-Cl, pH 7.5, 2.5 mM MgCl_2_, 0.5 mM CaCl_2_), supplemented with 20 U/ml RNase-free Recombinant DNase I (Invitrogen, Cat # AM2235) and incubated for 30 min at 37°C. RNA was then extracted with phenol-chloroform and twice precipitated with ethanol.

For RNA-seq analysis, two biological replicas were analyzed. Ribosomal RNA was removed from the samples using the Ribo-Zero rRNA depletion kit (Gram-negative bacteria; Illumina, # MRZGN126). Sequencing was done at the Laboratory for Molecular Biology and Cytometry Research at OUHSC. Whole transcriptome libraries were constructed using the Illumina TruSeq Total RNA V2 kit and established protocols. Raw data for each sample were mapped to the *P. aeruginosa* PAO1 genome for identification of genes expressed under each condition.

For qRT-PCR analysis, 2 μg of DNase treated total RNA was reverse transcribed into cDNA using the cDNA Reverse Transcription Kit (Invitrogen, Cat #: 4374966). PCR reactions were then set up using Power SYBR Green PCR Master Mix (Invitrogen, Cat #: 4367659), 1 μl of cDNA and 0.2 μM primers (Table S5). Real time PCR was conducted using ABI 7500 Real-Time PCR system (Applied Biosystems). 16S rRNA was used as an internal reference. Melting curve analysis confirmed that only one PCR product was generated in all cases. The relative gene expression was calculated using the 2-ΔΔC_t_ method (70). The data were averaged over at least 3 replica of cell cultures for 3 RT-PCR reactions each.

### Bioinformatic analysis

All statistical and bioinformatic analysis was conducted using the MATLAB R2017a platform. The abundance of each transcript was expressed as RPKM (Reads Per Kilobase per Million reads mapped), excluding the residual rRNA gene counts (less than 1.1% of total reads). Genes having zero reads in any of the samples (89 total) were removed from the analysis as well as *smc* and *mksB*. The false discovery rate (FDR) was calculated using the Storey correction (73). Hierarchical clustering and principle component analysis, PCA, were performed on the logarithms of the ratios of replica-averaged RPKM values found in mutants to those in the wild type strain. The pathway database compiled biological pathways from Kyoto Encyclopedia of Genes and Genomes (KEGG) (74), Virulence factor database (VFDB) (75), Pseudomonas Genome Database (PseudoCAP) (76, 77), and other literature sources (29, 33, 78-89). Venn diagrams were generated using MATLAB script written by Heil. Violin plots with overlapping boxplots were adapted from MATLAB script written by Bechtold (90).

## Acknowledgements

The authors thank the Laboratory for Molecular Biology and Cytometry Research at OUHSC for providing the Illumina MiSeq library construction, sequencing and analysis services for the RNAseq experiments. This work was supported by the award for the project number HR14-042 from Oklahoma Center for Advancement of Science and Technology to VVR. Our research is also supported in part by NIH Grants R21EY029015 and R21AI141927 (to VVR) R01EY024140 (to MCC), P30EY27125 (NIH CORE grant to Robert E. Anderson, OUHSC), a Presbyterian Health Foundation Equipment Grant (to Robert E. Anderson, OUHSC), and an unrestricted grant to the Dean A. McGee Eye Institute from Research to Prevent Blindness Inc. (http://www.rpbusa.org). We acknowledge Drs. Suzanne Fleiszig (University of California Berkeley) and Derek Royer (OUHSC) for helpful technical discussions. We also acknowledge Austin LaGrow (OUHSC) and the OUHSC Live Animal Imaging and Analysis Core facilities for technical assistance (P30EY27125), and Excalibur Pathology (Norman OK) for histology expertise. This work was presented in part at the 2016 Association for Research in Vision and Ophthalmology Meeting in Seattle WA.

## Legends to Supplemental materials

**Figure S1**. Hierarchical clustering of PAO1 condensin strains. (**A**) Heat map of expression changes. Genes are shown as rows and strains as columns. (**B**) The elbow method estimate of the number of significant clusters. The unexplained variance in the gene expression ratios was plotted against the postulated number of clusters (blue) and fitted to a bilinear interpolation (red).

**Figure S2**. (**A, B**) Doubling times of condensin mutants in M9+ medium missing (**A**) or containing (**B**) only the indicated amino acids, denoted using the single letter code. (**C, D**) Growth of *P. aeruginosa* PAO1 and its Δ*smc* variant in M9+ medium supplemented with the indicated amino acids.

**Table S1**. Composition of significant clusters

**Table S2**. Virulence regulators in significant clusters

**Table S3**. Significantly affected pathways in condensin mutants. Each worksheet shows the fold-change and false discovery rate (FDR) values for each gene included in a given pathway. The field Significance is marked Yes for genes with the expression level change greater than 2 and FDR < 0.1 in any of the mutants.

**Table S4**. Composition of the mixture of trace elements.

**Table S5**. Primers used for qRT-PCR analysis

